# Increasing pulse energy of 5Hz rTMS improves its efficacy in inducing excitatory aftereffects

**DOI:** 10.1101/652578

**Authors:** I Halawa, K Reichert, S Anil, M Sommer, W Paulus

## Abstract

**Introduction:** High frequency repetitive transcranial magnetic stimulation induces excitation when applied to the motor cortex as reflected by the increase MEP amplitudes after the stimulation. The effects differ according to pulse width, probably due to higher content of energy in the wider pulses and their ability to cause wider activation in comparison to shorter pulse shapes. Here we focus on the aftereffects generated with high frequency controllable pulse TMS (cTMS) with different pulse widths.

**Objectives:** To investigate the influence of pulse energy by using different stimulation intensities and pulse widths on the excitatory plastic aftereffects of high frequency (HF) rTMS.

**Methods:** Using a controllable pulse stimulator TMS (cTMS), we stimulated the hand motor cortex with 5 Hz rTMS applying 1200 bidirectional pulses with the main component widths of 80, 100 and 120 microseconds. 14 healthy subjects were initially investigated for six randomized sessions first with 80% RMT for anterior-posterior (AP) and posterior-anterior (PA). Then three more sessions using same pulse widths were added for 90% RMT anterior-posterior (AP).

**Results:** 80% HF rTMS did not produce any significant excitation in either AP or PA direction. 90% RMT AP stimulation with 100 and 120 microsecond-wide pulses were more excitatory, when compared to the 80 microsecond-wide pulses. We also found a correlation between the individual pulse energy and the plastic outcome of each session.

**Conclusions:** HF rTMS with wider pulses is more effective in producing excitatory aftereffects, an effect that correlated with the higher energy content of wider pulses and higher intensity.

**Significance:** The findings here suggest that wider pulses are capable of inducing more excitation, a fact that could contribute to better results in future clinical studies performed with wider pulses.

## Introduction

Therapeutic use of repetitive transcranial magnetic stimulation (rTMS) has been shown to have level A efficacy in the treatment of depression and chronic pain (Lefaucheur et al. 2014). The key rTMS parameters involved in its efficacy as stimulation frequency, intensity and number of pulses and sessions have been closely investigated (Rossini et al. 2015). However, there are few studies on the impact of pulse width on rTMS outcome because of the scarcity of devices with adjustable pulse widths (Peterchev et al. 2011).

The controllable pulse parameter TMS (cTMS3) device (Rogue research Inc., Montreal, Canada) allows to change the width of its near rectangular pulses using two capacitors, and two bipolar semiconductor transistors that alternate the current between the capacitors (Peterchev et al. 2014). Such customized pulses may be more efficient in delivering energy to the cortex (Goetz et al. 2013). With a rTMS frequency of 1 Hz we have already shown that only the widest pulse duration of 120μs leads to a significant increase in excitability, while the two shorter pulses shapes (40 and 80 Hz) produced significant inhibition (Halawa et al. 2019).

In an early LTP experiment, high frequency stimulation of rabbit cortex where pulse width were used as an analogue to intensity, stimulation with pulses longer than 100μs lead to significant potentiation of neurons (McNaughton et al. 1978). The authors also argued that cooperativity of multiple afferents had a comparable effect to increasing intensity and pulse widths of larger potentiation of the synapse.

Here, we used cTMS to test the effect of different pulse widths on the excitatory aftereffects of HF rTMS. In this study, we explored the interplay of coil orientation, pulse width, and pulse intensity on the aftereffects of a high frequency (5 Hz) rTMS train, a protocol well known to induce excitatory aftereffects (Hallett 2007; Ziemann et al. 2008).

## Methods

### Participants

We recruited fourteen healthy subjects, four males and ten females, mean age 23.5 ± 2.6 SD years. All participants were right-handed and free from any neurological or psychiatric disorders, took no centrally acting medications and had no contraindications to TMS. Resting motor threshold (RMT) above 70% MSO for a Magstim 200^2^ device was an exclusion criterion. That was done to avoid overheating of the cTMS coil delivering the rTMS specially for wider pulses.

We obtained written informed consent from each subject before participation. The local ethics committee of the University Medical Center Göttingen approved the study protocol, which conformed to the Declaration of Helsinki.

### Recordings

Motor evoked potentials (MEPs) were recorded from the first dorsal interosseous (FDI) muscle of the right hand with surface Ag–AgCl electrodes in a belly-tendon montage. The electromyography signals were amplified, band-pass filtered (2 Hz–2 kHz), and digitized at a sampling rate of 5 kHz with a micro-1401 AD converter (Cambridge Electronic Design Ltd., Cambridge, UK). All signals were stored digitally for offline analysis. The peak-to-peak MEP amplitude served as an index for M1 excitability. The participants were requested to relax the right FDI during the measurements. Individual traces contaminated by voluntary muscle contraction before the TMS pulse(s) were excluded from the analysis.

A Magstim 200^2^ (Magstim Co. Ltd., Whitland, UK) for measurement and a cTMS prototype 3 (cTMS3; Rogue Research Inc., Montreal, Canada) for intervention were used to deliver TMS over the M1 representation.

For the 80% AP and PA six and for the 90% AP condition three repeated, randomized sessions were performed and separated by at least one week to avoid carry-over effects. (fig 1):

**Figure 1:**
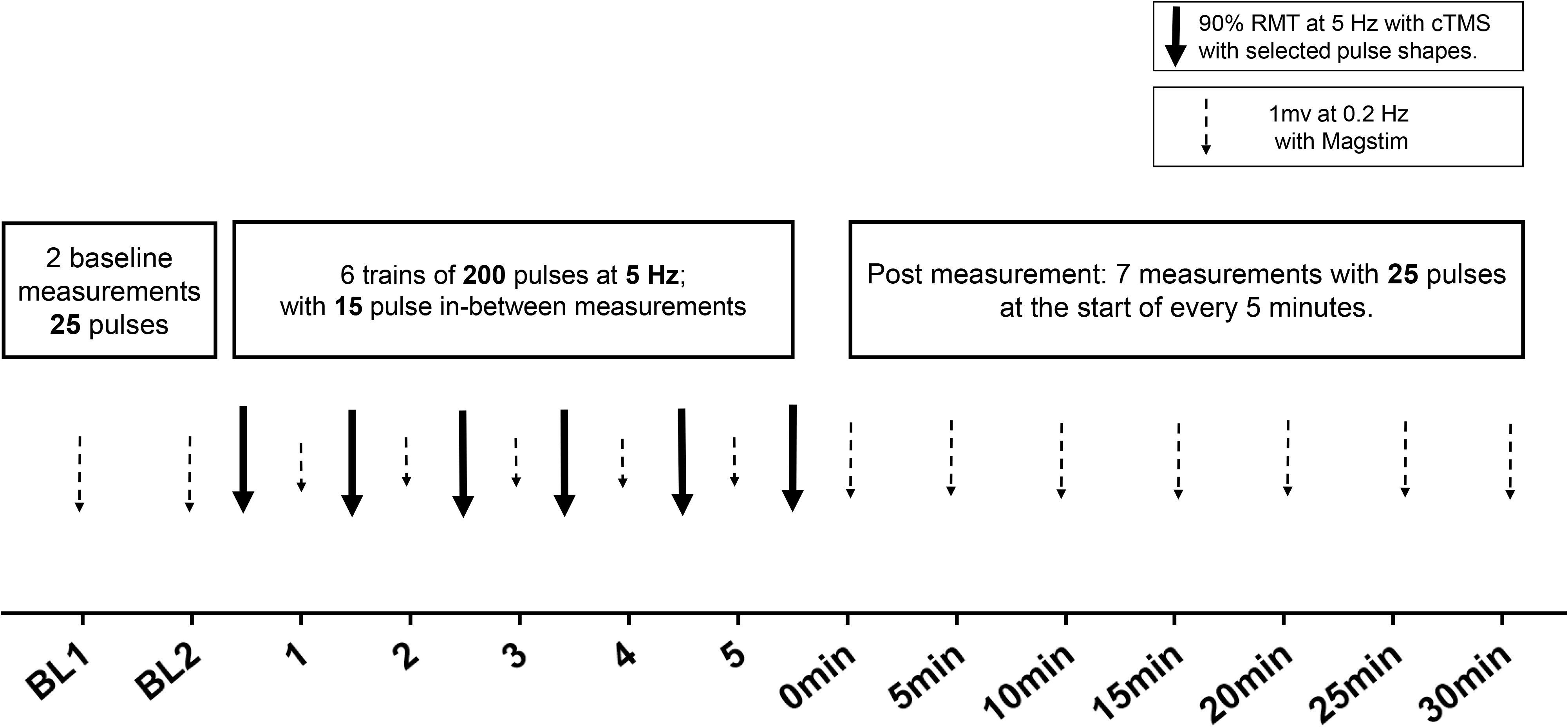
Diagram of the experiment timeline for each session of the experiment. Sessions were randomized and with at least one-week gaps to avoid possible carry over effects.

Step 1: Determining thresholds and baseline:

For each session, we used the Magstim 200^2^ with the D70 coil to determine the RMT and the MSO intensity that gave a response of approximately 1 mV for the baseline measurement intensity in the PA direction. In addition, we determined the RMT for the cTMS pulse shape being used for the intervention, both for the PA and the AP direction (coil rotated by 180 degrees), as a reference for the 5 Hz rTMS stimulation. The baseline measurements consisted of two 25-pulse measurements at 0.2 Hz with the previously determined MSO intensity giving a 1 mV response.

Step 2: The interventional cTMS stimulation:

As we could not use unidirectional pulses in high frequency stimulation as the capacitors of the device require a longer period of time to lose their charge (Peterchev et al. 2011), we used customized, balanced bidirectional pulse shapes which are easier to fire at higher frequencies. We used three widths of the main component (80μs, 100μs, and 120μs) in the PA and the AP direction at 80 % RMT and at 90 % for AP direction, always stimulating with 5 Hz (Fig 2).

**Figure 2:**
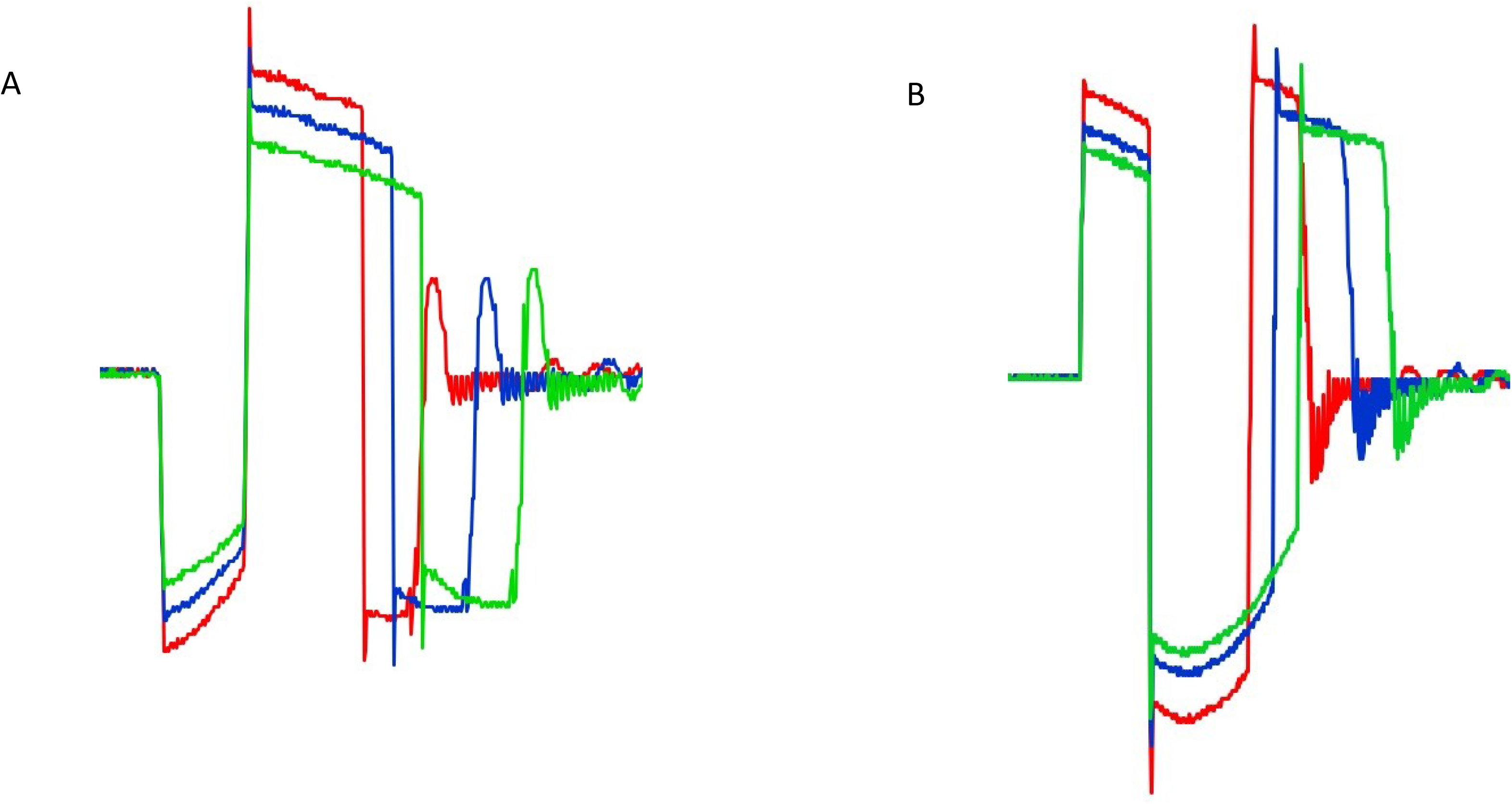
Pulse shapes used as detected by an external pickup coil and oscilloscope at their corresponding threshold intensities for: a) 5 Hz PA Stimulation. b)5 Hz AP Stimulation

1200 pulses were applied in six 200-pulse blocks separated by 15 MEP measurements at 0.2 Hz with the same intensity producing 1 mV determined in step 1 according to (Rothkegel et al. 2010).

Step 3: After the final rTMS pulse block, we recorded 25 pulses with the 1 mV intensity at a frequency of 0.2 Hz every five minutes for 30 minutes using the Magstim 200^2^.

#### Statistical analysis

We averaged the RMT values for each subject for each pulse shape. The effect of pulse width on RMT was analyzed using repeated measures ANOVA. For the MEP changes we used a multiple-way ANOVA for the pulse width, stimulation direction and evolution across time for baseline and after measurement time points. Then for each time point, we performed post-hoc multiple paired, two-tailed t-tests on the MEP amplitudes from the normalized 1 mV baseline for each condition with the baseline, using the Bonferroni-Dunn method to correct for multiple comparisons. We also averaged all post time points for the energy correlation. Pulse energy (U) was calculated for each stimulation intensity using the formula used by Peterchev and colleagues in their 2013 paper (Peterchev et al. 2013) 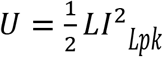 where I*Lpk* is the peak coil current, which we measured for the waveforms depicted in (Fig 2) with a single wire pick up coil.

## Results

### RMT

Wider pulse shapes had significantly lower RMTs (Fig 3.a) and higher energy contents than shorter pulse shapes (Fig 3.b) in the PA direction and the AP direction p < 0.0001 for both).

**Figure 3:**
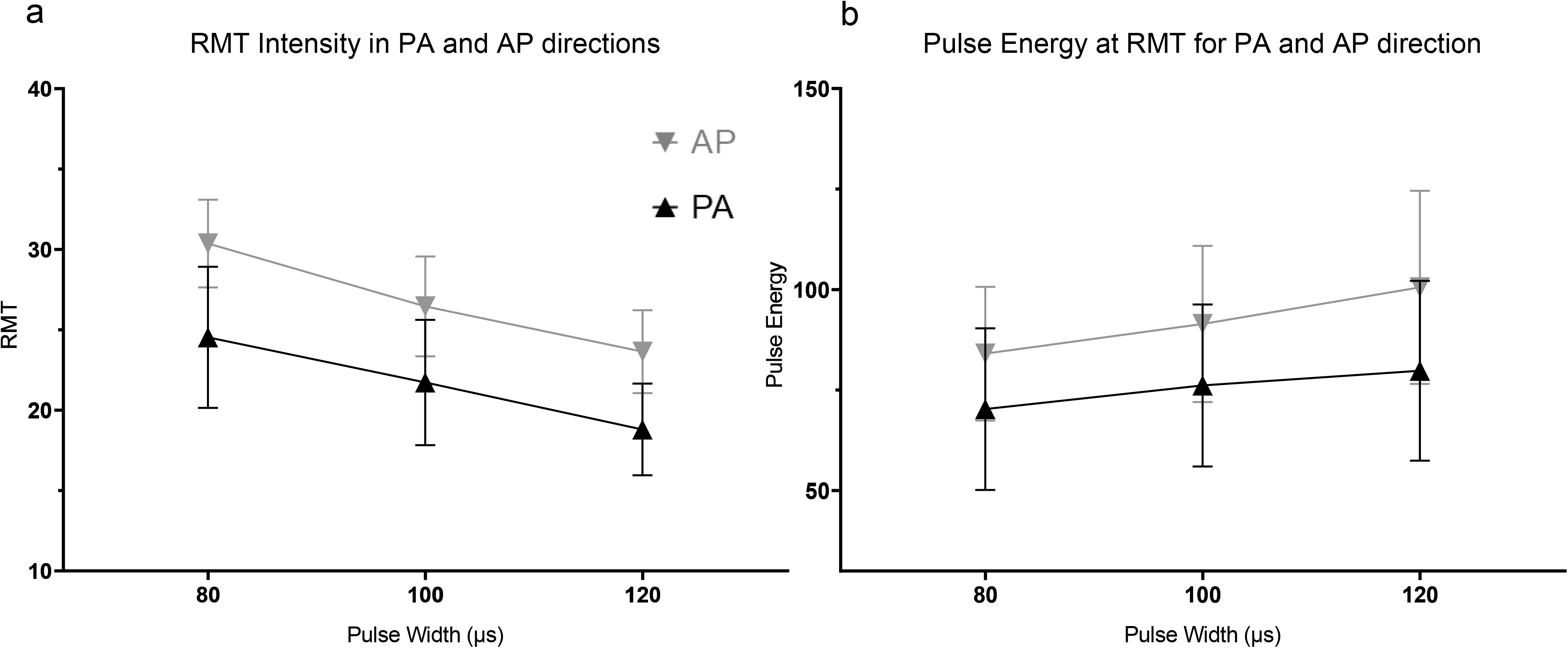
Effect of pulse width and stimulation direction on: a) Intensity thresholds b) Pulse Energy as determined in the methods section.

### Plastic aftereffects

We plotted the averaged normalized MEP data in the baseline phase, during stimulation and after stimulation (0 min to 30 min).

For the 80% RMT, PA directed 5 Hz rTMS, ANOVA revealed significant effect of varying pulse width P=0.0007, F=11.06 but no significant evolution across time points P=0.4857 and F=0.9913 (Fig.4).

**Figure 4:**
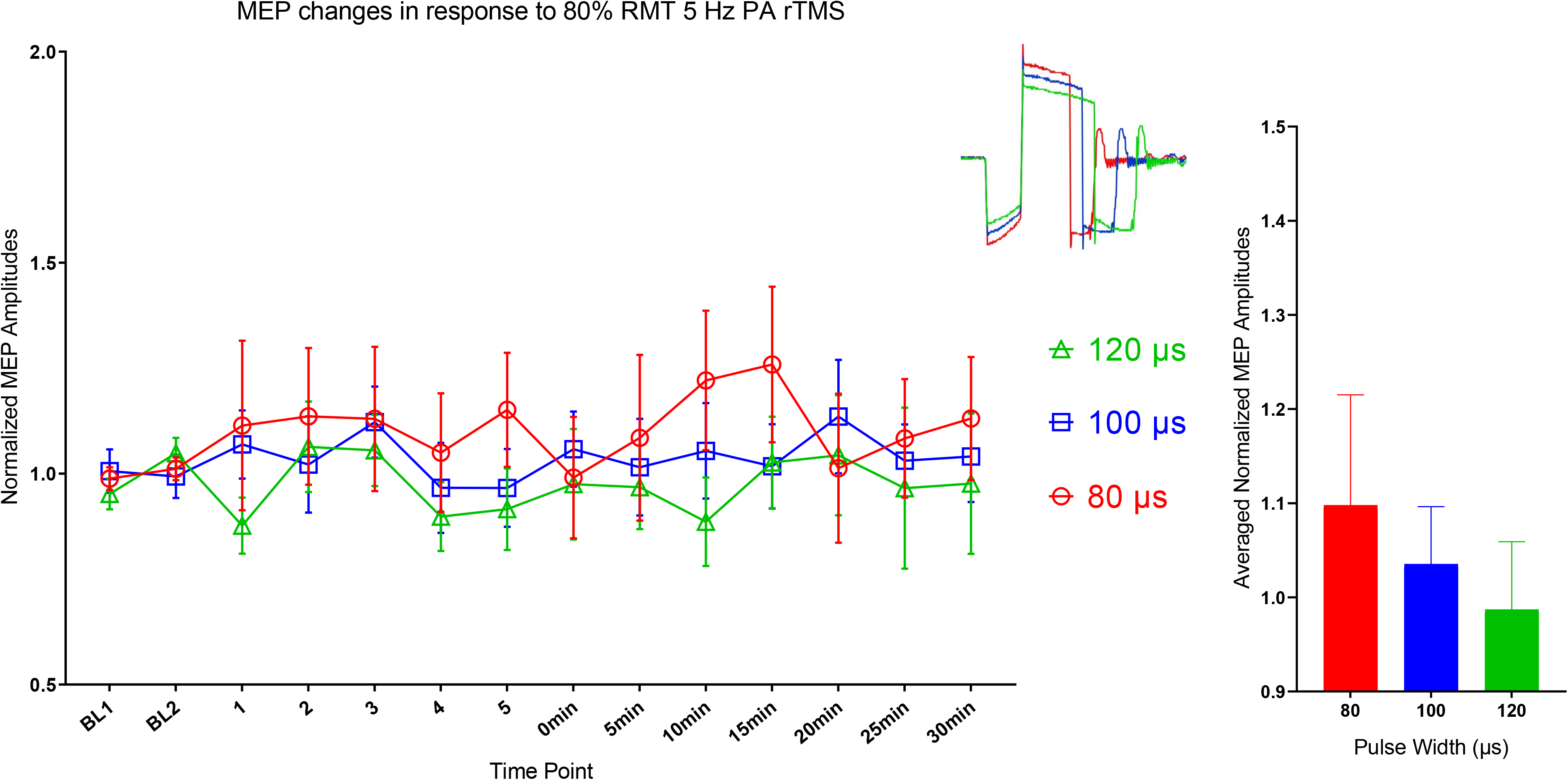
Aftereffects of 80% RMT 5 Hz stimulation using 80,100 and 120 μs main component in the PA direction. Pulse shapes used for stimulation are illustrated in corresponding colors in the top right panel.

For AP-directed stimulation also, there was a significant effect of pulse width change with p=0.0004; F=12.76.656 and no significant evolution across time points P=0.1791 and F=1.511 (Fig.5).

**Figure 5:**
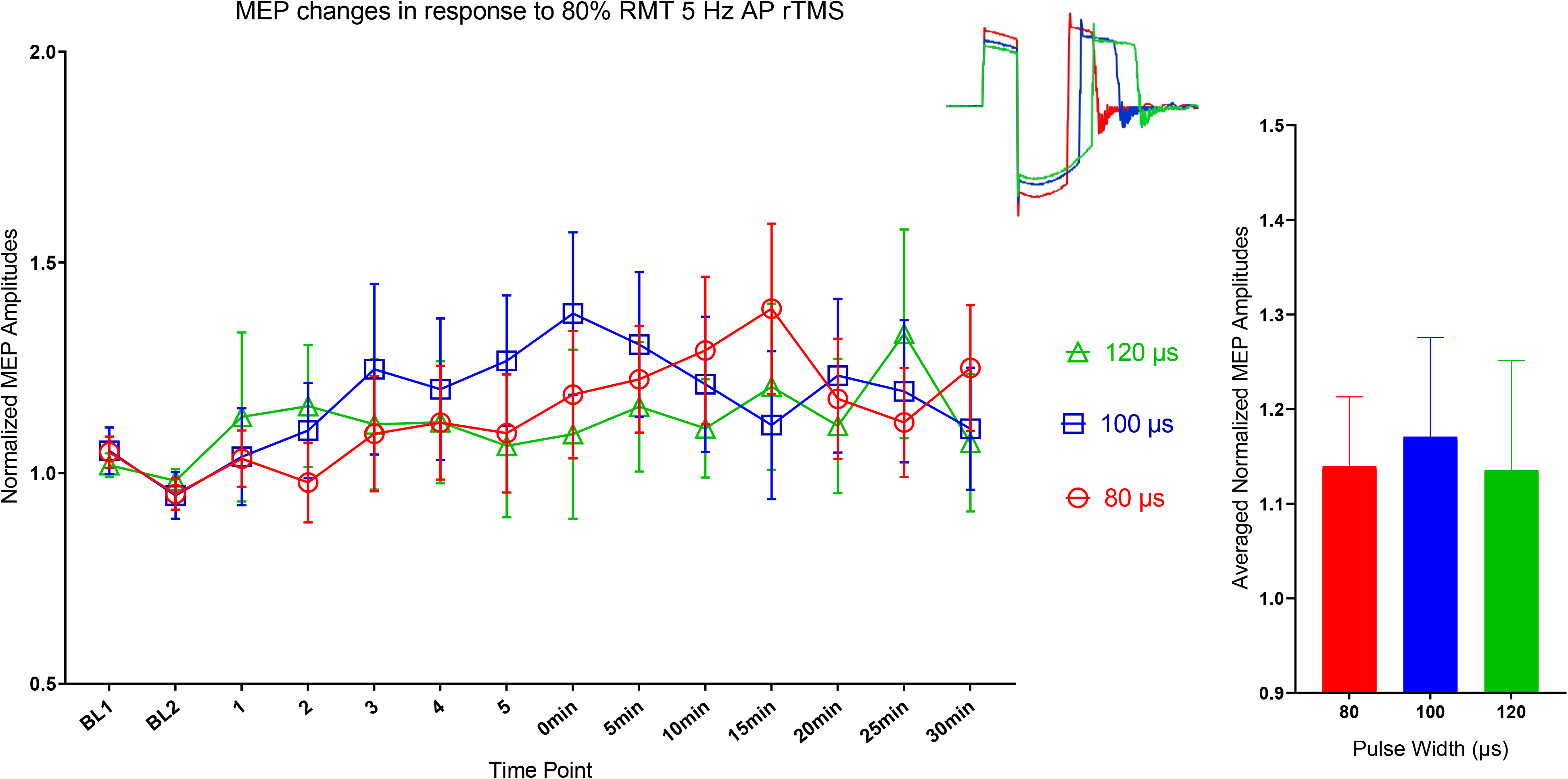
Aftereffects of 80% RMT 5 Hz stimulation using 80,100 and 120 μs main component in the AP direction. Pulse shapes used for stimulation are illustrated in corresponding colors in the top right panel.

Multiple corrected paired t-tests showed no significant difference across all time points. No significant difference was found in the baseline measurements.

For the 90% AP stimulation conditions, ANOVA revealed significant effects for pulse widths (p<0.0001; F= 18.23) and change over time (p=0.0017; F=3.356). Post-hoc t-tests showed that the 120 μs pulse shape produced significant excitation at six time points, the 100 μs condition produced excitation at four time points and the 80 μs condition exhibited a significant excitation at one time point (corrected p <0.009, paired two-tailed t-test, Fig.6). No significant difference was found within the baseline measurements.

**Figure 6:**
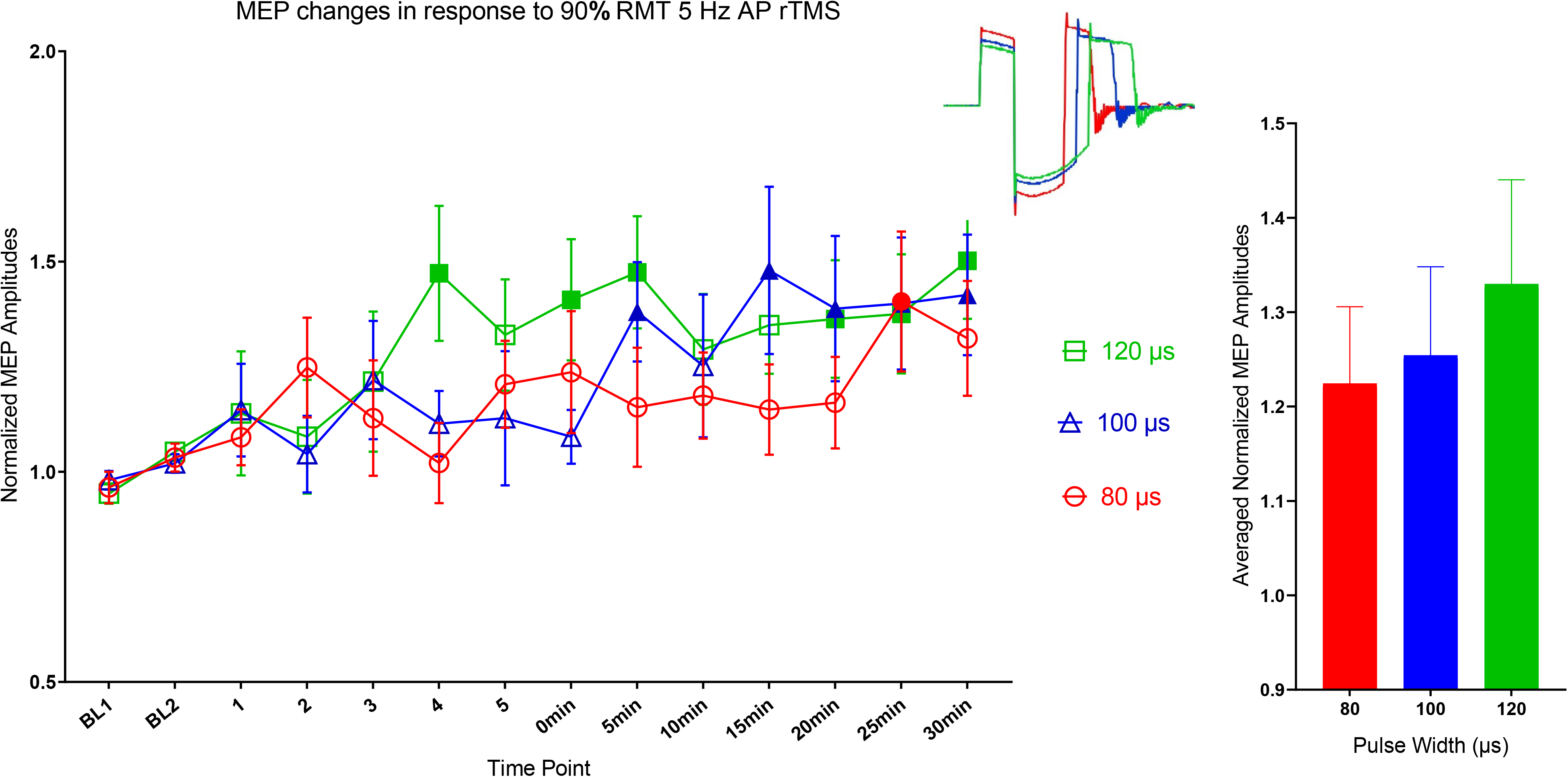
Aftereffects of 90% RMT 5 Hz stimulation using 80,100 and 120 μs main component in the AP direction. Pulse shapes used for stimulation are illustrated in corresponding colors in the top right panel. Solid shapes indicate significant shift from baseline.

When comparing the averaged change for all pulse widths, PA-directed stimulation did not produce any significant excitation (P=0.883, F=0.5083). The 80% AP condition exhibited significant shift from the baseline (P=0.45993, F=1.001). The 90% AP condition exhibited a significant shift from the baseline (P<0.0001, F=152.5) (Fig 7.a).

**Figure 7:**
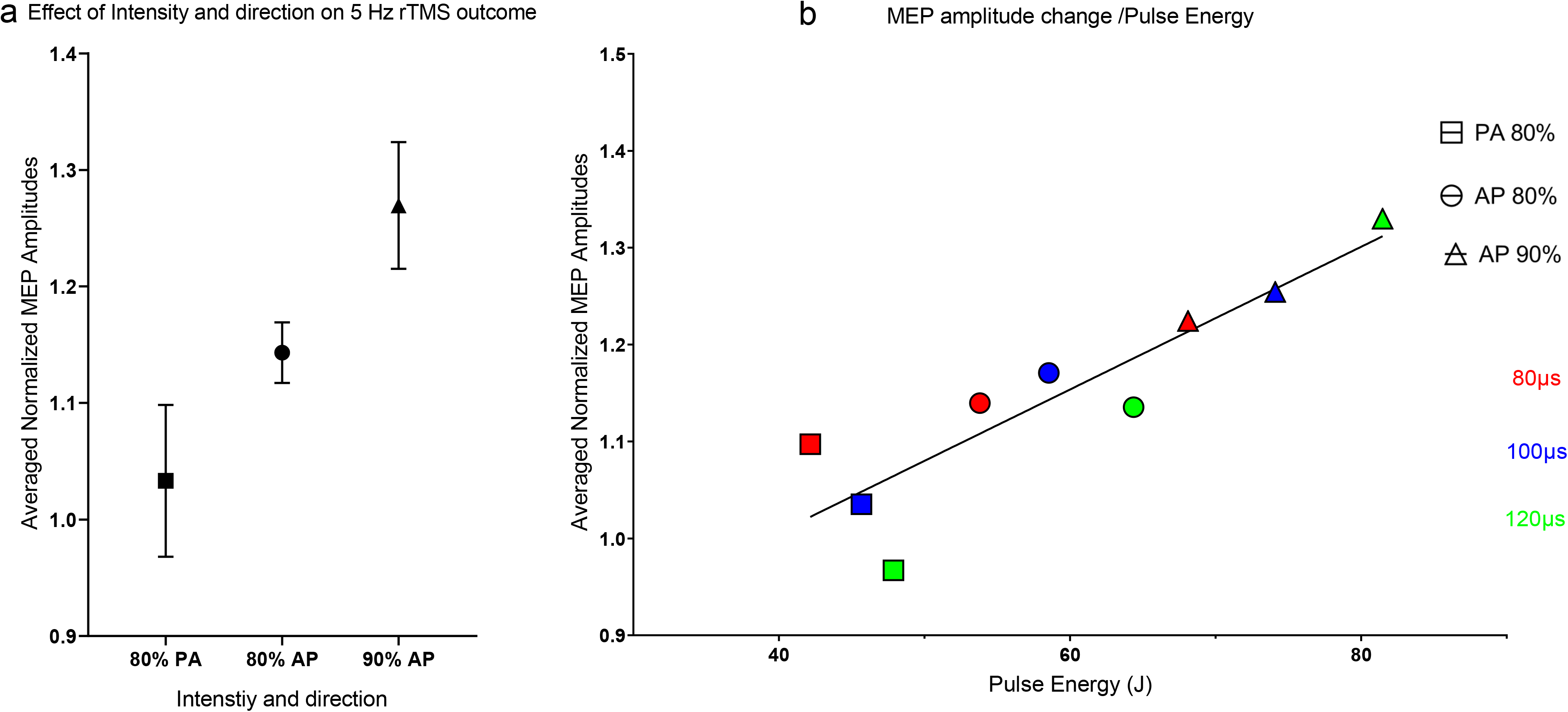
a.) Averaged Normalized MEP amplitudes in response for the 80% RMT PA, AP and 90% RMT AP 5 Hz stimulation. b.) Correlation between the averaged normalized MEP amplitudes in response to pulses with main component of 80, 100 and 120 μs for the 80% RMT PA, AP and 90% RMT with the corresponding energy content of the used pulse.

The averaged pulse energy content calculated for the corresponding stimulation intensity and pulse width exhibited a linear correlation with the averaged normalized MEP amplitudes with R^2^=0.79 (Fig 7.b).

## Discussion

In this study we showed that prolonging the pulse duration of HF rTMS leads to significant excitation for 90% RMT 5 Hz AP rTMS. Although there were no significant aftereffects after 80% RMT 5 Hz rTMS with pulse widths of 80, 100 and 120μs in either direction, we found that AP stimulation is more sensitive to pulse width change (p<0.001) and more effective when averaged in producing plastic aftereffects than PA stimulation (p<0.05). This has been demonstrated in many rTMS protocols especially for 5 Hz rTMS (Rothkegel et al. 2010; Sommer et al. 2013).

For the 90 % RMT AP-directed stimulation, the longest pulse with 120 μs main component produced the most significant excitation followed with the 100 μs. The shortest pulse shape with the 80 μs main component produced less excitation if compared to wider pulses, but still significant if compared to any of the 80% PA or AP conditions. The key finding then is that the 120 and 100 μs pulses with 5 Hz rTMS increased cortical excitability more than the 80μs pulses, and that with increasing the energy content of the pulse we got more excitation whether with increasing intensity or pulse widths.

In all subjects, RMT (Fig 3) decreased with increasing pulse width in agreement with previous reports (D’Ostilio et al. 2016). When looking at the correlation with total energy of the pulse as a function of the area of the pulse shape wider pulses require more energy to elicit responses in the cortex (Barker et al. 1991; Peterchev et al. 2013; Grill 2015). The higher total pulse energy could explain the seemingly greater efficacy of wider pulses when only pulse amplitude is considered (Modugno et al. 2001; Fitzgerald et al. 2006).

Neuronal models examining membrane properties for different parts of the neuron showed that because dendrites lack myelin (Aberra et al. 2018) and have a small diameter (Pashut et al. 2011), they respond preferentially to wider pulses (Rattay et al. 2012). That was also shown in an animal model experiment were short TMS pulses (~80μs) failed to produce any activation in L5 dendrites (Murphy et al. 2016). That implies that wider pulse shapes have a higher likelihood of stimulating dendrites probably because of their higher energy content. Dendritic involvement is also supported by the fact that more activation could be achieved by rotating magnetic fields or wider pulses (Rotem et al. 2014).

That effect could be explained by the larger dendritic capacitance mediated by their high resistance spines which allow them to passively amplify local synaptic depolarization up to 50-fold, where the higher spine neck resistance leads to increased cooperativity (Harnett et al. 2012). This capacitance was also shown by the fact that dendritic activation needed longer rTMS trains (i.e. more energy), but when they are eventually activated, they fired with higher amplitudes and for a longer period after stimulation ceases (Lee and Fried 2017). A similar effect was shown in single neurons where current injection into dendrites furthest from the soma produced longer and larger action potentials compared to somatic current injection (Larkum et al. 2007), especially in response to higher frequency and longer pulse width stimulation (Ledergerber and Larkum 2010).

Dendritic stimulation also produces back propagating potentials that would potentiate the anterograde potentials arising from somatic stimulation, thus producing LTP through associativity (Larkum 2004) or cooperativity and spike timing dependent plasticity (Lenz et al. 2015). This was supported by the fact that distal dendritic stimulation was more effective in producing cooperativity mediated LTP compared to proximal dendrite stimulation (Weber et al. 2016).

The significance of dendritic stimulation in producing lasting plastic aftereffects through LTP in response to rTMS is emphasized (Sjöström et al. 2008; Müller-Dahlhaus and Vlachos 2013). In this study we showed that increasing the pulse width increased the excitatory efficacy of HF rTMS. This is probably due to a stronger dendritic activation, resulting from their unique membrane properties and their important role in inducing synaptic plasticity.

Cooperativity mediated potentiation or excitation in TMS could be achieved by increasing energy content of individual pulses by increasing the intensity or the pulse widths to produce wider activation in the cortex.

## Acknowledgments

We thank the German Research Foundation (Deutsche Forschungsgemeinschaft, DFG) for funding the purchase of the cTMS device (DFG grant application: NST1525/18-1 FUGG). We also thank the German Academic Exchange Service (Deutscher Akademischer Austauschdienst, DAAD) and the Egyptian Ministry of Higher Education and Scientific Research (MHESR) for jointly funding the PhD scholarship for Islam Halawa.

We thank Prof. Thomas Crozier for language editing.

## References

Aberra AS, Peterchev AV, Grill WM. Biophysically Realistic Neuron Models for Simulation of Cortical Stimulation. 2018 Aug 20 [cited 2018 Nov 9]; Available from: http://biorxiv.org/lookup/doi/10.1101/328534

Barker AT, Garnham CW, Freeston IL. Magnetic nerve stimulation: the effect of waveform on efficiency, determination of neural membrane time constants and the measurement of stimulator output. Electroencephalogr Clin Neurophysiol Suppl. 1991;43:227–37.

D’Ostilio K, Goetz SM, Hannah R, Ciocca M, Chieffo R, Chen J-CA, et al. Effect of coil orientation on strength–duration time constant and I-wave activation with controllable pulse parameter transcranial magnetic stimulation. Clinical Neurophysiology. 2016 Jan;127(1):675–83.

Fitzgerald P, Fountain S, Daskalakis Z. A comprehensive review of the effects of rTMS on motor cortical excitability and inhibition. Clinical Neurophysiology. 2006 Dec;117(12):2584–96.

Goetz SM, Truong CN, Gerhofer MG, Peterchev AV, Herzog H-G, Weyh T. Analysis and Optimization of Pulse Dynamics for Magnetic Stimulation. Coles JA, editor. PLoS ONE. 2013 Mar 1;8(3):e55771.

Grill WM. Model-based analysis and design of waveforms for efficient neural stimulation. In: Progress in Brain Research [Internet]. Elsevier; 2015 [cited 2017 Oct 19]. p. 147–62. Available from: http://linkinghub.elsevier.com/retrieve/pii/S0079612315001375

Halawa I, Shirota Y, Neef A, Sommer M, Paulus W. Neuronal tuning: selective targeting of neuronal populations via manipulation of pulse width and directionality. Brain Stimulation. 2019 Apr;S1935861X19302049.

Hallett M. Transcranial Magnetic Stimulation: A Primer. Neuron. 2007 Jul;55(2):187–99.

Harnett MT, Makara JK, Spruston N, Kath WL, Magee JC. Synaptic amplification by dendritic spines enhances input cooperativity. Nature. 2012 Nov;491(7425):599–602.

Larkum ME. Top-down Dendritic Input Increases the Gain of Layer 5 Pyramidal Neurons. Cerebral Cortex. 2004 Apr 27;14(10):1059–70.

Larkum ME, Waters J, Sakmann B, Helmchen F. Dendritic Spikes in Apical Dendrites of Neocortical Layer 2/3 Pyramidal Neurons. Journal of Neuroscience. 2007 Aug 22;27(34):8999–9008.

Ledergerber D, Larkum ME. Properties of Layer 6 Pyramidal Neuron Apical Dendrites. Journal of Neuroscience. 2010 Sep 29;30(39):13031–44.

Lee SW, Fried SI. Enhanced Control of Cortical Pyramidal Neurons With Micromagnetic Stimulation. IEEE Transactions on Neural Systems and Rehabilitation Engineering. 2017 Sep;25(9):1375–86.

Lefaucheur J-P, André-Obadia N, Antal A, Ayache SS, Baeken C, Benninger DH, et al. Evidence-based guidelines on the therapeutic use of repetitive transcranial magnetic stimulation (rTMS). Clinical Neurophysiology. 2014;125(11):2150–2206.

Lenz M, Platschek S, Priesemann V, Becker D, Willems LM, Ziemann U, et al. Repetitive magnetic stimulation induces plasticity of excitatory postsynapses on proximal dendrites of cultured mouse CA1 pyramidal neurons. Brain Struct Funct. 2015 Nov;220(6):3323–37.

McNaughton BL, Douglas RM, Goddard GV. Synaptic enhancement in fascia dentata: Cooperativity among coactive afferents. Brain Research. 1978 Nov;157(2):277–93.

Modugno N, Nakamura Y, MacKinnon C, Filipovic S, Bestmann S, Berardelli A, et al. Motor cortex excitability following short trains of repetitive magnetic stimuli. Exp Brain Res. 2001 Oct;140(4):453–9.

Müller-Dahlhaus F, Vlachos A. Unraveling the cellular and molecular mechanisms of repetitive magnetic stimulation. Front Mol Neurosci [Internet]. 2013 [cited 2019 Jun 2];6. Available from: http://journal.frontiersin.org/article/10.3389/fnmol.2013.00050/abstract

Murphy SC, Palmer LM, Nyffeler T, Müri RM, Larkum ME. Transcranial magnetic stimulation (TMS) inhibits cortical dendrites. Elife. 2016;5:e13598.

Pashut T, Wolfus S, Friedman A, Lavidor M, Bar-Gad I, Yeshurun Y, et al. Mechanisms of Magnetic Stimulation of Central Nervous System Neurons. Graham LJ, editor. PLoS Computational Biology. 2011 Mar 24;7(3):e1002022.

Peterchev AV, DʼOstilio K, Rothwell JC, Murphy DL. Controllable pulse parameter transcranial magnetic stimulator with enhanced circuit topology and pulse shaping. Journal of Neural Engineering. 2014 Oct 1;11(5):056023.

Peterchev AV, Goetz SM, Westin GG, Luber B, Lisanby SH. Pulse width dependence of motor threshold and input–output curve characterized with controllable pulse parameter transcranial magnetic stimulation. Clinical Neurophysiology. 2013 Jul;124(7):1364–72.

Peterchev AV, Murphy DL, Lisanby SH. Repetitive transcranial magnetic stimulator with controllable pulse parameters. Journal of Neural Engineering. 2011 Jun 1;8(3):036016.

Rattay F, Paredes LP, Leao RN. Strength–duration relationship for intra- versus extracellular stimulation with microelectrodes. Neuroscience. 2012 Jul;214:1–13.

Rossini PM, Burke D, Chen R, Cohen LG, Daskalakis Z, Di Iorio R, et al. Non-invasive electrical and magnetic stimulation of the brain, spinal cord, roots and peripheral nerves: Basic principles and procedures for routine clinical and research application. An updated report from an I.F.C.N. Committee. Clinical Neurophysiology. 2015 Jun;126(6):1071–107.

Rotem A, Neef A, Neef NE, Agudelo-Toro A, Rakhmilevitch D, Paulus W, et al. Solving the Orientation Specific Constraints in Transcranial Magnetic Stimulation by Rotating Fields. PLOS ONE. 2014 May 2;9(2):e86794.

Rothkegel H, Sommer M, Paulus W. Breaks during 5Hz rTMS are essential for facilitatory after effects. Clinical Neurophysiology. 2010 Mar;121(3):426–30.

Sjöström PJ, Rancz EA, Roth A, Häusser M. Dendritic Excitability and Synaptic Plasticity. Physiological Reviews. 2008 Apr;88(2):769–840.

Sommer M, Norden C, Schmack L, Rothkegel H, Lang N, Paulus W. Opposite Optimal Current Flow Directions for Induction of Neuroplasticity and Excitation Threshold in the Human Motor Cortex. Brain Stimulation. 2013 May;6(3):363–70.

Weber JP, Andrásfalvy BK, Polito M, Magó Á, Ujfalussy BB, Makara JK. Location-dependent synaptic plasticity rules by dendritic spine cooperativity. Nat Commun. 2016 Sep;7(1):11380.

Ziemann U, Paulus W, Nitsche MA, Pascual-Leone A, Byblow WD, Berardelli A, et al. Consensus: Motor cortex plasticity protocols. Brain Stimulation. 2008 Jul;1(3):164–82.

